# ASXL1 Directs Neutrophilic Differentiation via Modulation of MYC and RNA Polymerase II

**DOI:** 10.1101/2020.09.14.295295

**Authors:** Theodore P. Braun, Joseph Estabrook, Lucie Darmusey, Daniel J. Coleman, Zachary Schonrock, Brittany M. Smith, Akram Taherinasab, Trevor Enright, Cody Coblentz, William Yashar, Rowan Callahan, Hisham Mohammed, Brian J. Druker, Theresa A. Lusardi, Julia E. Maxson

## Abstract

Mutations in the gene Additional Sex-Combs Like 1 (ASXL1) are recurrent in myeloid malignancies as well as the pre-malignant condition clonal hematopoiesis, where they are universally associated with poor prognosis. An epigenetic regulator, ASXL1 canonically directs the deposition of H3K27me3 via the polycomb repressive complex 2. However, its precise role in myeloid lineage maturation is incompletely described. We utilized single cell RNA sequencing (scRNA-seq) on a murine model of hematopoietic-specific ASXL1 deletion and identified a specific role for ASXL1 in terminal granulocyte maturation. Terminal maturation is accompanied by down regulation of Myc expression and cell cycle exit. ASXL1 deletion leads to hyperactivation of Myc in granulocyte precursors and a quantitative decrease in neutrophil production. This failure of normal developmentally-associated Myc suppression is not accompanied by significant changes in the landscape of covalent histone modifications including H3K27me3. Examining the genome-wide localization of ASXL1 in myeloid progenitors revealed strong co-localization with RNA Polymerase II (RNAPII) at the promoters and spread across the gene bodies of transcriptionally active genes. ASXL1 deletion results in a decrease in RNAPII promoter-proximal pausing in granulocyte progenitors, indicative of a global increase in productive transcription, consistent with the known role of ASXL1 as a mediator of RNAPII pause release. These results suggest that ASXL1 inhibits productive transcription in granulocyte progenitors, identifying a new role for this epigenetic regulator and highlighting a novel potential oncogenic mechanism for ASXL1 mutations in myeloid malignancies.

## Introduction

Recurrent mutations in the epigenetic regulator Additional Sex-Combs Like 1 (ASXL1) are associated with universally poor prognosis in myeloid malignancy (Gelsi-Boyer et al., 2012; Kim et al., 2017; Pratcorona et al., 2012; Tefferi et al., 2014). Furthermore, mutations in ASXL1 also occur in the premalignant condition clonal hematopoiesis, where they are associated with a high rate of progression to myeloid leukemia (Abelson et al., 2018; Genovese et al., 2014; Jaiswal et al., 2014). In spite of this, little is known about the role of ASXL1 in myeloid lineage development, which is significant barrier to the development of therapeutic approaches to combat this mutation.

Asx family members were originally described in drosophila, where deletion results in both anterior and posterior homeotic phenotypes in the same individual, suggesting that that ASXL1 may regulate chromatin structure to facilitate both activation and repression of homeotic genes (Milne et al., 1999). In mammals, there are three related ASXL family members, of which only ASXL1 and ASXL2 are expressed in adult hematopoietic cells. Prior work has suggested that ASXL1, in particular, directs the core polycomb repressive complex 2 (PRC2) to the Hoxa locus, leading to deposition of repressive histone 3 lysine 27 trimethylation (H3K27me3) (Abdel-Wahab et al., 2012, 2013). Indeed, knockdown of ASXL1 in leukemia cell lines results in a loss of H3K27me3 at the Hoxa locus and upregulation of Hox genes, in particular Hoxa9. Hox genes play an important role in hematopoietic stem/progenitor self-renewal and are downregulated during maturation (Alharbi et al., 2013; Pineault et al., 2002). Aberrant expression of HOXA9 is an important driver of leukemia maintenance and thus aberrant Hox gene expression has been implicated as an important mediator of ASXL1 mutation-dependent biology (Ayton and Cleary, 2003; Bullinger et al., 2004).

Initially characterized as loss of function, recent work has attributed gain of function properties to mutant ASXL1, suggesting the possibility of neomorphic oncogenic activity (Abdel-Wahab et al., 2012, 2013; Inoue et al., 2013; Nagase et al., 2018; Yang et al., 2018). Multiple murine models of ASXL1 mutation have been developed, all displaying hematopoietic phenotypes, but each with differing features. Further complicating the understanding of ASXL1 mutations is an incomplete understanding of the lineage specific functions of this global epigenetic regulator. We therefore set out to understand the role of ASXL1 in normal myeloid development.

Recent advances in single cell analysis have revealed that many epigenetic regulatory factors play specific roles in lineage development that are relatively unique to particular cell subsets (Izzo et al., 2020; Viny et al., 2019). In order to understand the normal biological role of ASXL1 in normal hematopoiesis, we utilized scRNA sequencing and CUT&Tag-based low-input epigenetic profiling to investigate the specific role of ASXL1 in myeloid cell development. These studies revealed that ASXL1 is required for terminal granulocyte production via regulation of global transcription in the committed neutrophil progenitor. Our studies demonstrate little variance in key covalent histone modifications, instead suggesting a role for ASXL1 in modulating the activity of RNA Polymerase II. These findings provide an important basis for the investigation of the impact of mutant ASXL1 on hematopoiesis, identifying key cell types and biological processes that are uniquely dependent on ASXL1 function.

## Results

### Single Cell RNA Sequencing Defines Multiple Lineages in Murine Bone Marrow

To assess the impact of ASXL1 deletion on hematopoietic lineage outputs, we performed single cell RNA sequencing on lineage negative (Lin-) hematopoietic progenitor cells from MX1 Cre ASXL1^FL/FL^ or littermate control Cre negative mice (henceforth referred to as ASXL1^Δ/Δ^ and ASXL1^WT^ respectively) (Figure 1A). We examined two time points after induction of recombination via Poly I:C administration: 4 weeks and greater than 6 months. At the former timepoint, peripheral blood counts are normal, while they begin to decline in mice >6 months of age. In total, we profiled 53,579 cells, with data integration revealing 25 transcriptionally distinct subsets. Immature progenitor cell populations were associated with the expression of genes such as Cdk6 whereas distinct downstream lineages were defined by the expression of specific gene subsets (Figure 1 B, C, Supplementary Figure 1A). Erythroid lineage cells demonstrated high expression of hemoglobin subunits and carbonic anhydrase (Car1), while lymphoid cells expressed immunoglobulin subunits and cytokine receptors (i.e. Jchain in B cells and Il2rb in T/NK cells). Myeloid lineage cells demonstrate an immature myeloid progenitor population (IMP) from which both granulocytic and monocytic cells arise, delineated by the differential expression of the lineage defining transcription factors Cebpe and IRF8 respectively.

**Figure 1.**
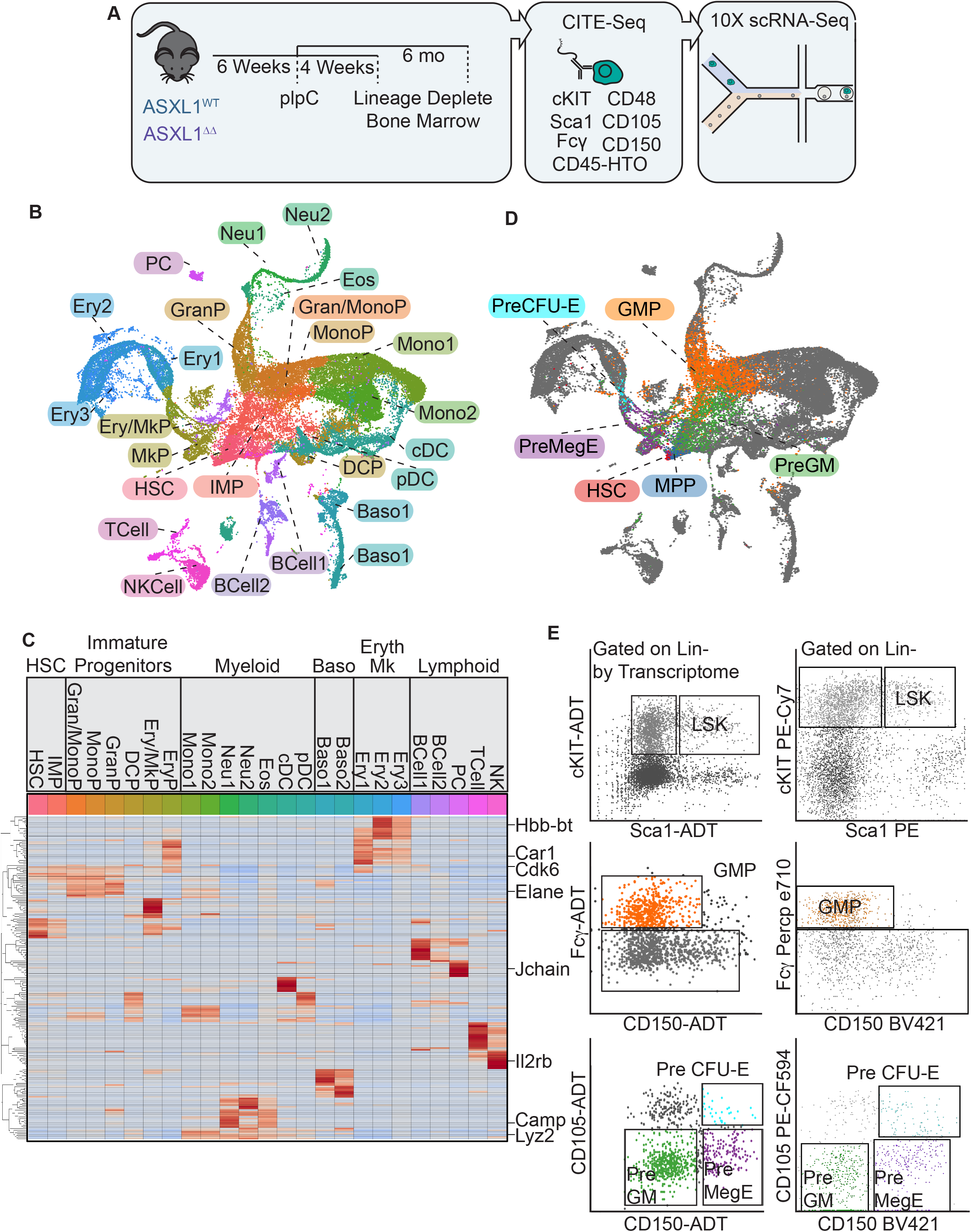
Combined Transcriptomic and Proteomic Profiling Reveals Multi-Lineage Maturation Through Known Stem and Progenitor Populations. **A**. Bone marrow was isolated from ASXL1^WT^ and ASXL1 ^Δ/Δ^ mice 4 weeks and 6 months after induction of recombination with Poly I:C (n=3-4/group). Lineage positive cells were depleted using immunomagnetic purification and labeled with the indicated CITE-seq antibodies. Each individual mouse was labeled with a unique anti CD45 HTO antibody allowing for mouse-level bioinformatic deconvolution. Single cell transcriptional profiling was then performed using the Chromium platform (10X Genomics). **B**. UMAP projection demonstrating transcriptionally defined clusters identified using published datasets and data integration. **C**. Marker genes for transcriptionally defined clusters. **D**. UMAP projection demonstrating clusters defined by CITE-Seq. **E**. Gating strategy for CITE-seq pseudo flow and for parallel flow cytometry.

To confirm the transcriptional identities assigned to cluster, we performed Cellular Indexing of Transcriptomes and Epitopes by Sequencing (CITE-seq) for key surface markers associated with hematopoietic stem and progenitor cell identity (Stoeckius et al., 2017). CITE-seq identified long term hematopoietic stem cells (HSCs), Multipotent Progenitor Cells (MPPs), Pre Granulocyte-Macrophage Progenitor (PreGM), Granulocyte Macrophage Progenitors (GMPs), Pre Megakaryocyte Erythroid Progenitors (PreMegE), and Pre Colony Forming Unit Erythroid (Pre CFU-E) within the transcriptionally defined clusters in a manner highly analogous to flow cytometry performed in parallel (Figure 1D, E) (Oguro et al., 2013; Pronk et al., 2007). Indeed, CITE-seq defined clusters mapped to the expected transcriptionally defined clusters with a high degree of concordance (Supplementary Figure 1B). Collectively, these results establish a baseline map of the cellular constituents and lineages comprising murine bone marrow and represent a platform to investigate the role of ASXL1 in cell fate determination.

### ASXL1 Deletion Results in an Early Loss of Bone Marrow Granulocytic Progenitors

ASXL1^Δ/Δ^ mice are known to develop peripheral neutropenia when they reach >6 months of age, however at earlier timepoints they maintain peripheral blood neutrophil counts within the normal range (Abdel-Wahab et al., 2013). In contrast, scRNA-seq of bone marrow progenitors reveals a marked defect in the output of the granulocyte lineage 4 weeks after the induction of recombination that worsens with ageing (Figure 2A, B). In contrast, the majority of other lineages demonstrate largely intact cell numbers. ASXL1^Δ/Δ^ mice maintain normal granulocyte maturation until they reach the Neu1 transcriptionally defined cluster, shortly after exiting the CITE-seq defined GMP cluster. Lineage tracing analysis placing all cells along maturation trajectory and assigning pseudo-time values, reveals uniform cell counts in all clusters prior to the committed neutrophils, confirming that the defect in maturation occurs at the post-GMP stage (Figure 2C, D). To further confirm the granulocyte lineage defect associated with ASXL1 deletion, we examined the capacity of ASXL1-deficient bone marrow to produce mature neutrophils in ex-vivo culture (Supplementary Figure 2A). This demonstrated that neutrophil progenitors can be produced in normal numbers from ASXL1^Δ/Δ^ bone marrow, but that these progenitors fail to differentiate in normal numbers upon withdrawal of stemness-promoting cytokines. Finally, although at baseline ASXL1^Δ/Δ^ mice demonstrate largely normal peripheral blood neutrophil counts, they demonstrate reduced peripheral granulocytosis upon in vivo GCSF challenge (Supplementary Figure 2B).

**Figure 2.**
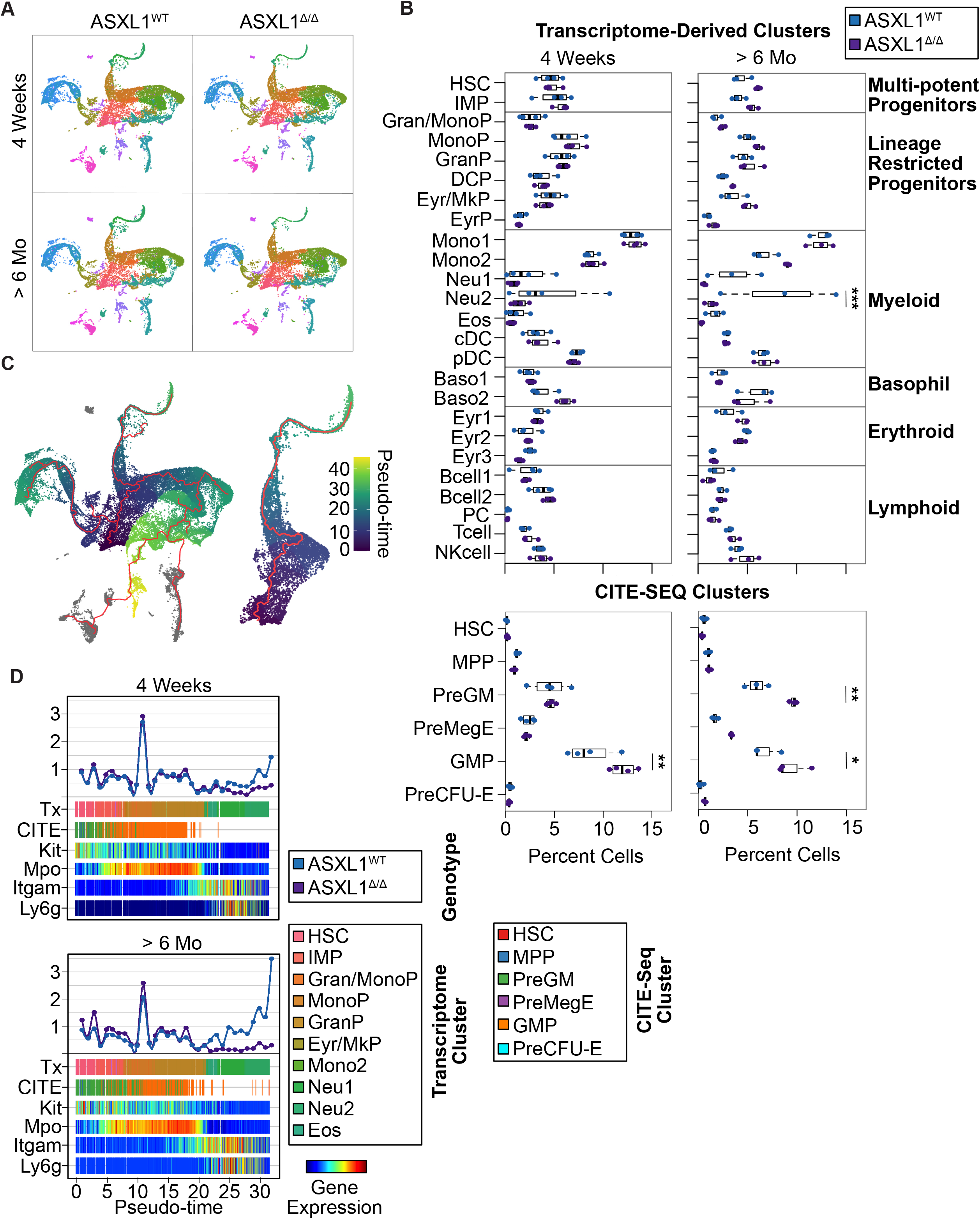
Deletion of ASXL1 Leads to a Loss of Granulocyte Maturation Potential. **A**. UMAP projections for ASXL1^WT^ and ASXL1 ^Δ/Δ^ at 4 weeks and 6 months post induction of recombination (n=3-4/group). **B**. Cell number by cluster within transcriptome and CITE-seq derived clusters. **C**. Cell trajectories for all lineages and expanded view of granulocytic lineage. **D**. Linearized granulocyte trajectory showing transcriptional and CITE-seq clusters along with expression of key marker genes. In all cases plots indicate mean and standard error. * = p<0.05, ** =p< 0.01, *** =p<0.001 by two-way ANOVA with Tukey HSD posttest.

To further elucidate the nature of the maturation defect, we performed CytoTRACE analysis, which predicts maturation by correlation of maturity with the total number of genes expressed per cell (Gulati et al., 2020). This revealed that transcript heterogeneity increases as cells leave the long-term stem cell pool and reaches a maxima at points of fate decision, then falling again as cells approach terminal differentiation (Figure 3A, B). Examining the CytoTRACE score along the granulocytic lineage and integrating this information with transcriptionally defined clusters, we found that the maturation trajectory through the granulocyte precursors (GranP) bifurcates between ASXL1^Δ/Δ^ and ASXL1^WT^ cells, with the pathway taken by the ASXL1 deficient cells ultimately being productive of fewer maturing neutrophils (Figure 3C, D). These data collectively demonstrate that disruption of ASXL1 results in an early defect in granulocyte lineage maturation.

**Figure 3.**
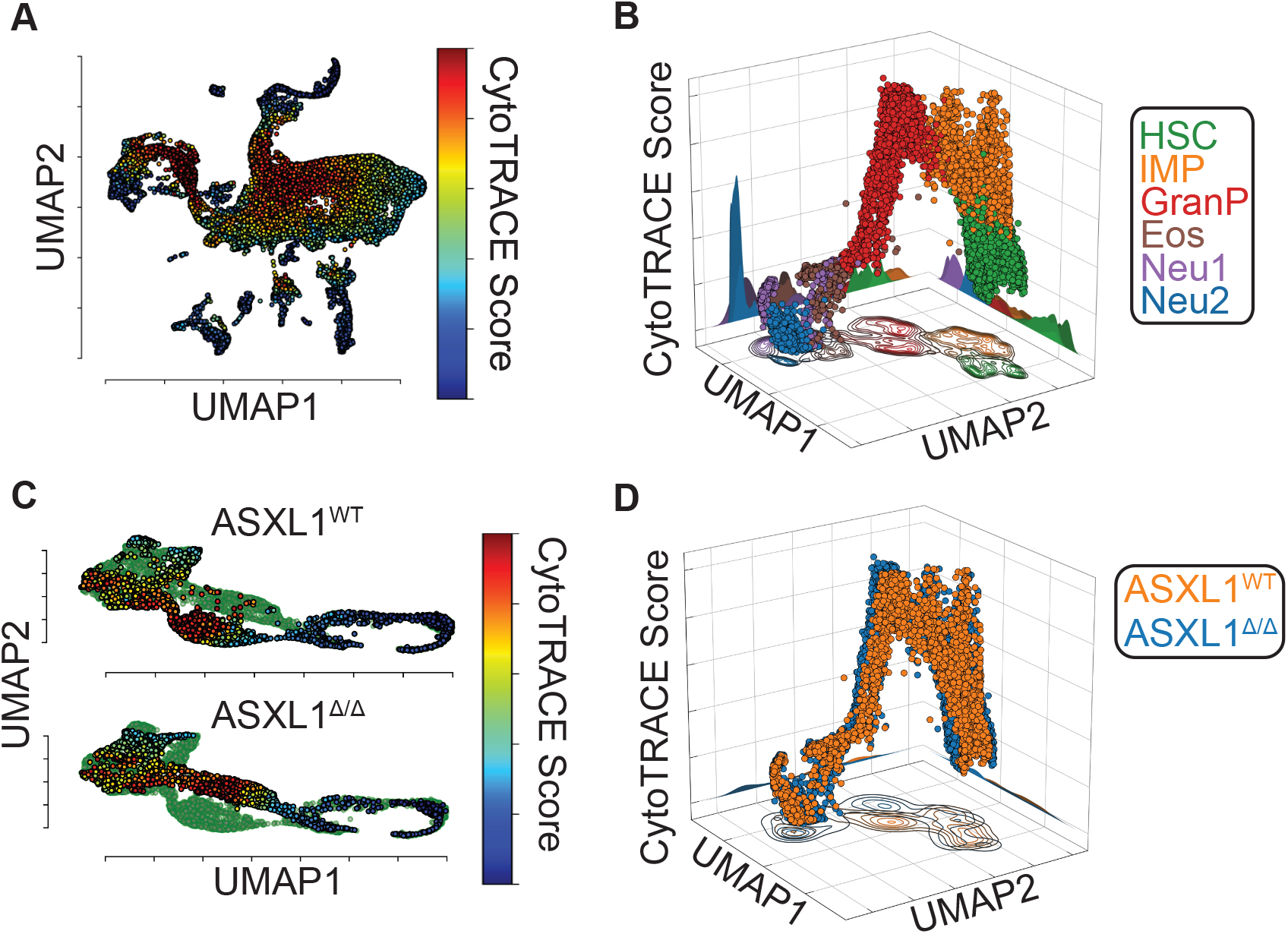
Deletion of ASXL1 Results in a Transcriptionally Divergent Maturation Pathway. **A**. UMAP projection of CytoTRACE scores. **B**. Three-dimensional projection of granulocyte lineage maturation with transcriptional UMAPs on X and Y axis and Cyto-TRACE score on Z axis and projections against Z-axis indicating cell pileup. **C**. Split UMAPs showing granulocyte lineage maturation in ASXL1^WT^ and ASXL1 ^Δ/Δ^ mice. **D**. Same projection as **B** showing granulocyte lineage maturation in ASXL1^WT^ and ASXL1 ^Δ/Δ^ mice.

### ASXL1 Deletion Results in Perturbation of the Myc Network in Maturing Granulocytes

To assess the transcriptional impacts of ASXL1 deletion on granulocyte maturation, we investigated differential gene expression down the granulocyte trajectory. At both early and late time points, the number of differential genes detected increased with maturation reaching a maximum in the GranP cluster and then falling with further maturation (Supplementary Figure 3A). Gene ontology analysis revealed enrichment for terms associated with cell growth, RNA splicing and chromatin organization upregulated in ASXL1^Δ/Δ^ cells while terms associated with immune function were down regulated (Supplementary Figure 3B). To elucidate the dynamics of transcriptional networks that occur in response to ASXL1 deletion, we performed a network analysis using CausalPath (Figure 4A) (Babur et al., 2018). Globally, this analysis revealed a similar pattern to that seen with the differential expression analysis. Network complexity increased with maturation up to the point of the GranP cluster, again decreasing markedly thereafter (Figure 4B). These networks nucleated in the HSC cluster with a Myb-dependent network, expanding with maturation to involve Myc-associated factor X (MAX), NFkB, and the anti-apoptotic family member Bcl2 in the IMP population (Figure 4C) In the GranP cluster, the network then expanded to include Myc and cell cycle regulators E2f3, Ccnd2 and Mcm6 among others (Figure 4D). Crucially, exit from the GranP population is associated with cell cycle arrest, and these findings argue that the loss of granulocyte maturation is associated with dysregulation of this Myc-dependent developmentally regulated process (Supplementary Figure 3C).

**Figure 4.**
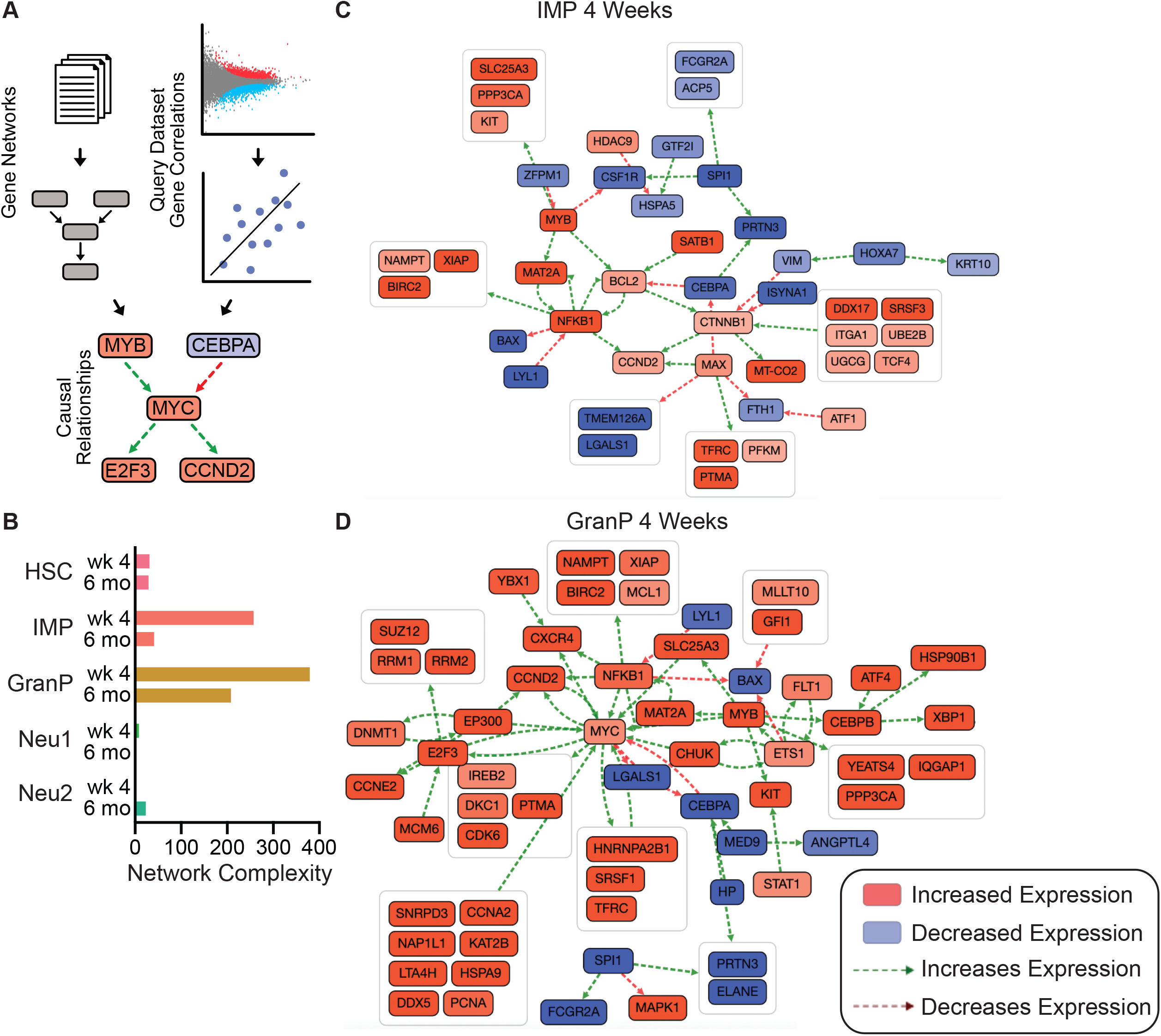
Deletion of ASXL1 Leads to MYC Network Activation in Granulocyte Progenitors. **A**. Schematic of CausalPath analysis which utilizes literature derived gene regulatory networks and gene correlations from the query dataset to predict causal relationships. **B**. Number of network edges as a measure transcriptional network complexity between ASXL1^WT^ and ASXL1 ^Δ/Δ^ mice. **C**. Differential network activation in IMP population 4 weeks post induction of recombination. **D**. Differential network activation in granulocyte progenitor population 4 weeks post induction of recombination.

### ASXL1 Regulates Polycomb Repressive Complex 2-mediated Gene Repression in Myeloid but not Granulocytic Progenitors

Prior work, predominantly in immature cell populations, has extensively characterized the role of ASXL1 in regulating PRC2 mediated deposition of repressive H3K27me3 histone modifications (Abdel-Wahab et al., 2012, 2013). To characterize the dynamics of this chromatin modification during granulocyte maturation, we developed a flow cytometric sorting strategy to purify myeloid progenitors, granulocytic progenitors and mature bone marrow neutrophils. We then utilized Cleavage Under Targets and Tagmentation (CUT&Tag) based genome wide profiling of H3K27me3 across the trajectory of granulocyte maturation (Figure 5A, B) (Kaya-Okur et al., 2019). To characterize the differences between genotypes, we performed a differential signal enrichment analysis between ASXL1^Δ/Δ^ and ASXL1^WT^ mice in each cell type. This analysis revealed the largest number of differential H3K27me3 peaks in myeloid progenitors with few identified in granulocyte progenitors (Figure 5C). Gene ontology analysis of these differentially marked regions in myeloid progenitors revealed enrichment of terms associated RNA polymerase activity and multiple neuronal specific ion channels, consistent with a role for ASXL1 in mediating PRC2-dependent repression of neuronal phenotypes in immature hematopoietic cells (Supplementary Figure 4A). We also assessed the impact of ASXL1 deletion on regions of H3K27me3 that are gained or lost with granulocytic maturation. This revealed that maturation was associated with largely normal deposition of H3K27me3 in ASXL1 deficient cells (Supplementary Figure 4B-D). Consistent with this, global comparison of signal tracks between ASXL1^Δ/Δ^ and ASXL1^WT^ cells revealed strong correlation in neutrophil precursors and mature neutrophils but weaker correlation in myeloid progenitors (Figure 5D). In addition, within these sorted cell populations, ASXL1 deletion failed to markedly change the total amount of H3K27me3, which peaked in abundance in the granulocyte precursor population (Figure 5E),

**Figure 5.**
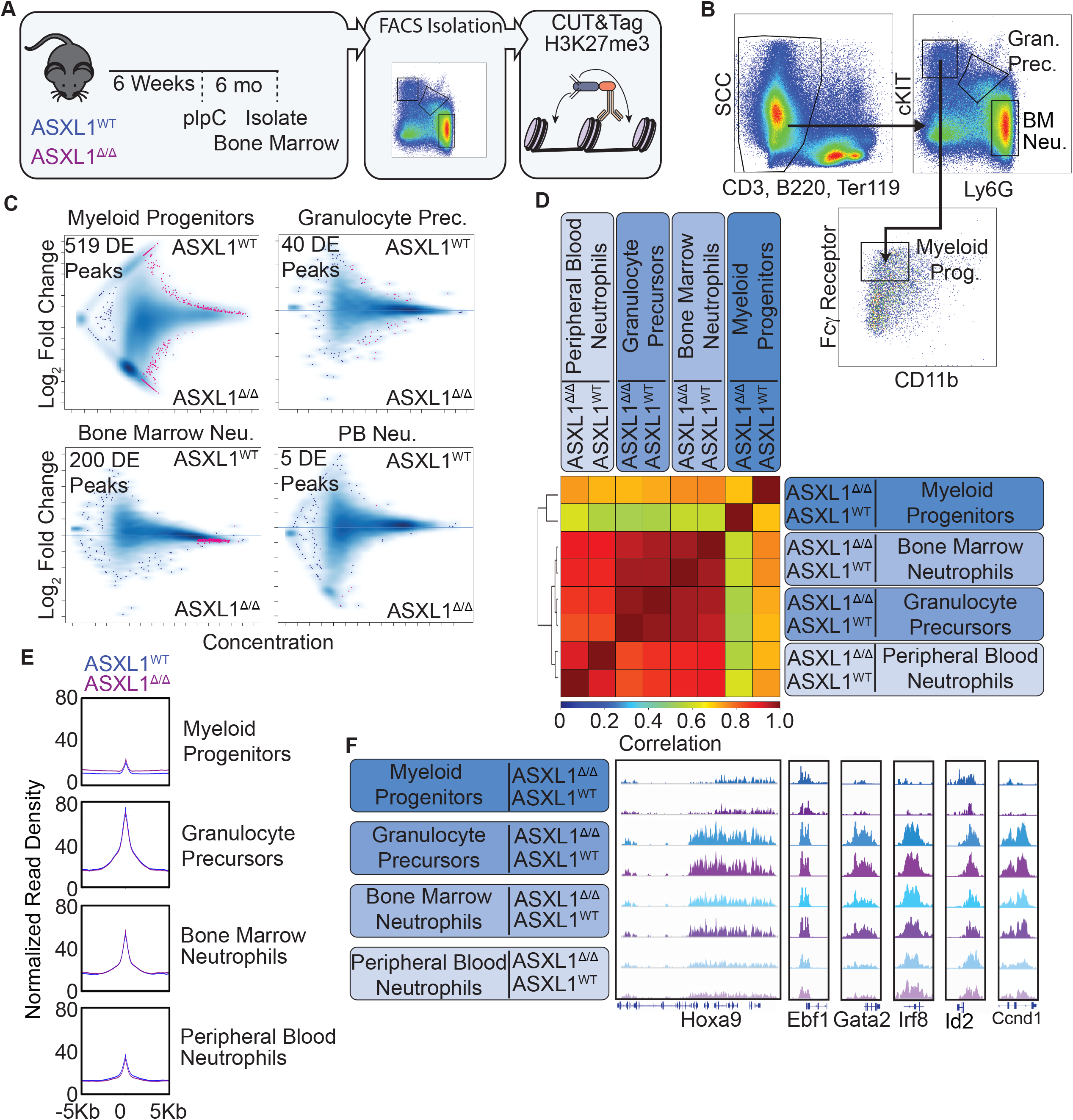
ASXL1 Deletion Produces Minimal Impacts on H3K27me3 Deposition in Granulocyte Progenitors. **A**. Experimental schematic. Bone marrow from ASXL1^WT^ and ASXL1 ^Δ/Δ^ mice was harvested 6 months after induction of recombination via poly I:C administration, maturing granulocyte lineage cells isolated by FACS, and subjected to CUT&Tag for H3K27me3. **B**. Example flow sorting strategy for Myeloid Progenitors, Gran-ulocyte Progenitors and Bone Marrow Neutrophils. **C**. MA plots for differential enrichment analysis of H3K27me3 in Myeloid Progenitors, Granulocyte Progenitors, Bone Marrow Neutrophils and Peripheral Blood Neutrophils. Red dots indicate regions of differential H3K27me3 enrichment. **D**. Spearman Correlation of Genome-wide H3K27me3 signal in indicated bone marrow sub-populations. **E**. Mean normalized read density at H3K27me3 peaks in indicated bone marrow sub populations. **F**. Representative tracks of H3K27me3 signal in indicated bone marrow sub-populations.

Despite these differences, PRC2 activity appears largely unaffected by ASXL1 deletion. Normal hematopoiesis is associated with downregulation of Hoxa9 expression. Consistent with this, we observed increased H3K27me3 signal at the Hox locus with differentiation and heterochromatic spread to cover Hoxa9 that was equivalent in both ASXL1^Δ/Δ^ and ASXL1^WT^ cells (Figure 5F). Furthermore, appropriate repression of transcription factors associated with differential lineage fates occurred normally in ASXL1^Δ/Δ^ mice as represented by Ebf1 (Lymphoid), Gata2 (erythroid) and Irf8 (monocyte). Even for regions identified as differentially marked in the myeloid progenitor such as Id2 and Ccnd1, differentiation down the granulocytic lineage was associated with marked gains in H3K27me3 at these loci, overcoming any small differences present at the earlier stage. Collectively these data reveal that granulocytic differentiation is associated with the deposition of repressive H3K27me3 marks and that while some modest differences are noted in ASXL1^Δ/Δ^ cells (particularly early progenitors), the vast majority of deposition of H3K27me3 associated with normal myeloid maturation occurs in an ASXL1 independent manner. Critically, minimal differences in H3K27me3 are noted in neutrophil precursors, the cell type with the greatest transcriptional dysregulation upon ASXL1-deletion, arguing that these differential gene expression changes occur through alternate mechanisms.

### Prom1/CD133 Marks Committed Granulocyte Precursors and Serves as a Target for Purification

Our scRNA-seq analysis revealed that committed granulocyte precursors demonstrate the greatest transcriptional perturbation upon deletion of ASXL1 (Figure 2). However, profiling of the H3K27me3 reveals that this repressive mark is most dynamic in less mature myeloid progenitors and displays little variance upon ASXL1 deletion in the more committed granulocyte precursors (Figure 5). To better characterize this population, we searched our scRNA seq dataset for lineage-specific expression of surface markers that were not differentially expressed upon ASXL1 deletion. We identified Prom1 which encodes the surface marker CD133 and is expressed exclusively on granulocyte progenitors (Figure 6A). Therefore, we developed an immunomagnetic purification strategy to isolate CD133 positive cells. This isolation successfully purified cKit-positive, Ly6G-dim cells from murine bone marrow with minimal contamination from either cKit-negative cells or Ly6G-negative cells (Figure 6B). To confirm that these cells also demonstrated similar transcriptional differences to those observed in the scRNA-seq dataset, we performed bulk RNA seq on ASXL1^Δ/Δ^ and ASXL1^WT^ CD133-positive granulocyte progenitors. This analysis similarly revealed multiple low-level transcriptional perturbations with a preponderance of upregulated ribosomal proteins (Figure 6C). Furthermore, GO analysis revealed an enrichment for Myc targets in genes upregulated after ASXL1 deletion (Figure 6D). Thus, CD133 purification isolates granulocyte progenitors with a similar transcriptional profile to those identified in scRNA sequencing from lineage depleted marrow.

**Figure 6.**
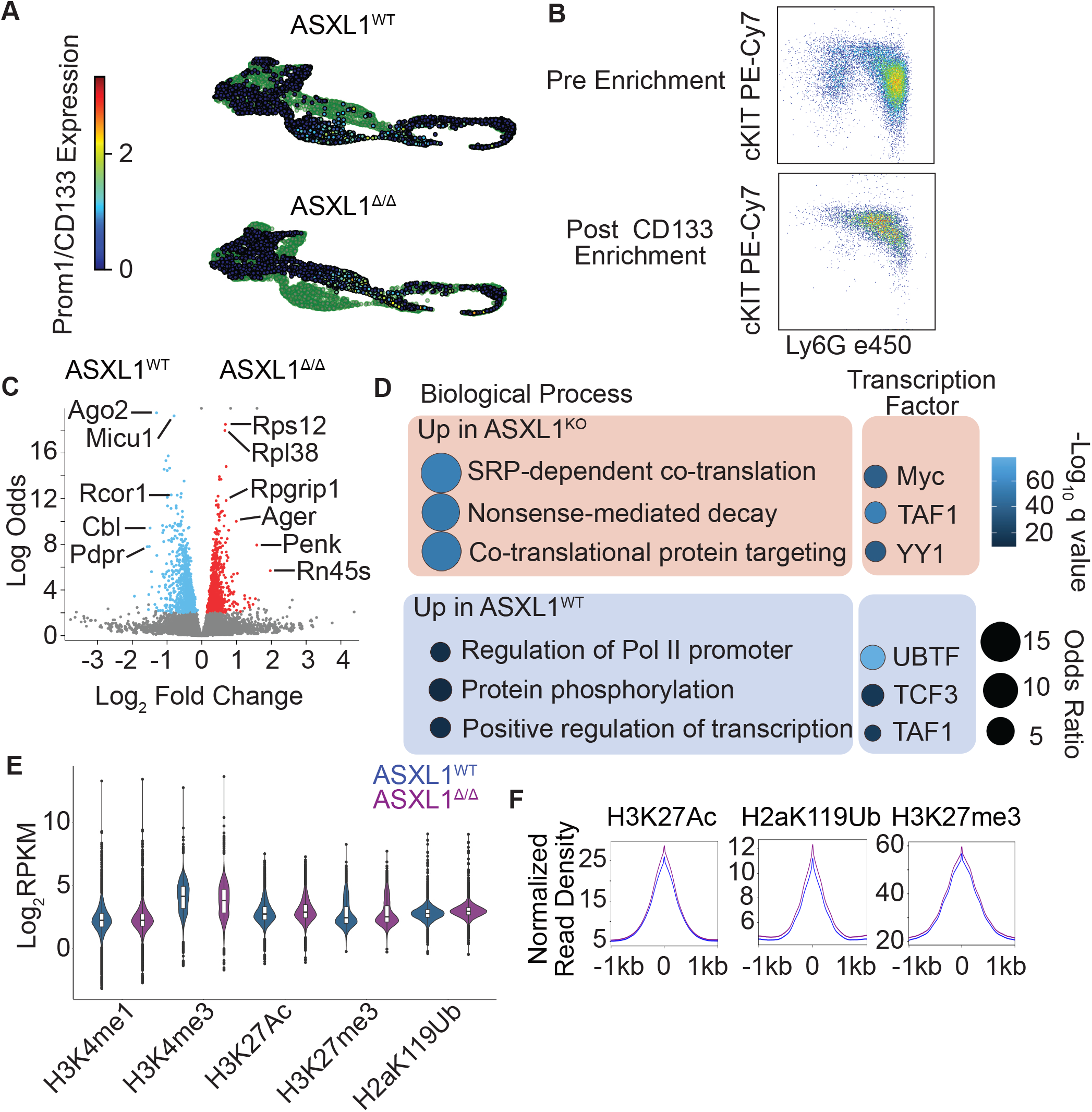
Granulocytic Progenitors Show Minimal Perturbation of the Chromatin Landscape in Response to ASXL1 Deletion. **A**. UMAP projection of Prom1/CD133 in granulocyte progenitors. **B**. Representative FACS plots pre and post CD133-immuno-magnetic purification. **C**. Volcano plot from bulk RNA-seq on purified granulocyte progenitors from ASXL1^WT^ and ASXL1 ^Δ/Δ^ mice (n=4/group). Red dots represent significantly upregulated genes **D**. GO analysis on bulk RNA-seq from **C**. with top 3 GO biological process and Encode/ChIA consensus transcription factors shown. **E**. Total signal in peaks for each covalent histone modification assessed by CUT&Tag in CD133 positive granulocyte progenitors from ASXL1^WT^ and ASXL1 ^Δ/Δ^ (n=2-4/group). **F**. Mean signal in peaks for indicated histone marks in CD133 positive granulocyte progenitors from ASXL1^WT^ and ASXL1 ^Δ/Δ^ mice (n=4/group).

### Comprehensive Chromatin State Profiling of Granulocytic Progenitors Reveals Minimal Perturbations in Response to ASXL1 Deletion

Our data suggest that H3K27me3 marks are relatively static in flow cytometrically isolated granulocytic progenitors (Figure 5), arguing that ASXL1 may modulate transcriptional output via alternate epigenetic mechanisms. We therefore performed CUT&Tag for multiple covalent histone modifications enabling the global segmentation of chromatin into distinct functional domains. Specifically, we examined H3K4me1 which is broadly associated with enhancers, H3K4me3 which is present at active promoters, H3K27Ac which is present at active enhancers and promoters. In addition, we assessed H3K27me3 to correlate our results with our prior findings in sorted cells. Finally, we profiled H2aK119Ub which is associated with PRC1-mediated gene repression and can specifically be altered by ASXL1 via activation of the BAP1 deubiquitinase (Asada et al., 2018). Globally, the total abundance of all marks was not substantially altered by ASXL1 deletion (Figure 6E, F), nor were substantial numbers of differential peaks detected between ASXL1^WT^ and ASXL1^Δ/Δ^ cells (Supplementary Figure 5A).

To determine whether specific chromatin states might be altered by ASXL1 deletion, we then utilized a hidden Markov model-based approach to segment the genome into 8 distinct chromatin states, identifying promoters, enhancers, regions of activation and repression (Supplemental Figure 5B, C). Globally, there were minimal shifts in repressed chromatin states, and greater, but still subtle changes in regions of active chromatin (Supplementary Figure 5D). Specifically, the greatest degree of chromatin remodeling was observed at enhancers, with exchange of regions between activate and primed states. To investigate the relationship between each mark and gene expression, we examined the correlation between changes in histone mark read counts and RNA read counts. Globally, no strong correlation was observed between changes in either H3K27Ac and H2aK119ub and gene expression (Supplementary Figure 5E, F). These data collectively demonstrate that ASXL1 deletion leads to relatively minimal changes in the epigenetic landscape with stronger changes observed in activating rather than repressive chromatin marks. However, none of these changes clearly correlate with differential gene expression arguing that ASXL1 directly impacts transcription via alternate mechanisms.

### ASXL1 Localizes to Regions of Active Transcription and Regulates RNA-Polymerase II Activity

Given the strong Myc-dependent gene expression signature we observed in scRNA seq and bulk RNA seq datasets as well as the role in Myc in regulating RNAPII pause release, we hypothesized that ASXL1 might be modifying gene expression through direct regulation of RNAPII (Eberhardy and Farnham, 2002; Price, 2010). We therefore compared ASXL1 ChIP-seq signal from cKIT+ murine bone marrow progenitors with covalent histone modifications profiled in CD133 positive cells (Nagase et al., 2018). In addition, we profiled RNAPolII binding genome wide in CD133+ neutrophil progenitors. Examining global signal, we saw the greatest degree of correlation between ASXL1 and RNAPolII signal and the least correlation with H3K27me3 (Figure 7A). These findings were confirmed in an alternate human cell type using published ChIP-seq data (HEK293 cells, Supplementary Figure 6). Indeed, ASXL1 was largely distributed at the transcriptional start sites of actively transcribed genes (for example the myeloid transcription factor Cebpa, Figure 7B). To assess the activity of RNAPolII in ASXL1^KO^ granulocytic progenitors, we examined the RNAPII pause index in which a ratio of proximal promoter read density is divided by the read density across the remainder of the gene body (Figure 7C) (Nagase et al., 2018). We observed that ASXL1^Δ/Δ^ granulocyte progenitors had consistently lower pause indices than ASXL1^WT^ cells (7D-F). This occurred predominantly at genes with high promoter proximal PolII signal suggestive of PolII pausing. At individual loci, this occurred in multiple patterns with PolII signal progressing down the gene body (i.e. Rpl37), decreased signal at the proximal promoter (i.e. Cebpa) or increased signal at the point of PolII termination (i.e. Eef2, Figure 7G). These results are consistent with the known physical interaction of ASXL1 with PolII in non-hematopoietic cell types and argue for ASXL1 playing a role in directly modulating the activity of RNAPolII (Zhang et al., 2018).

**Figure 7.**
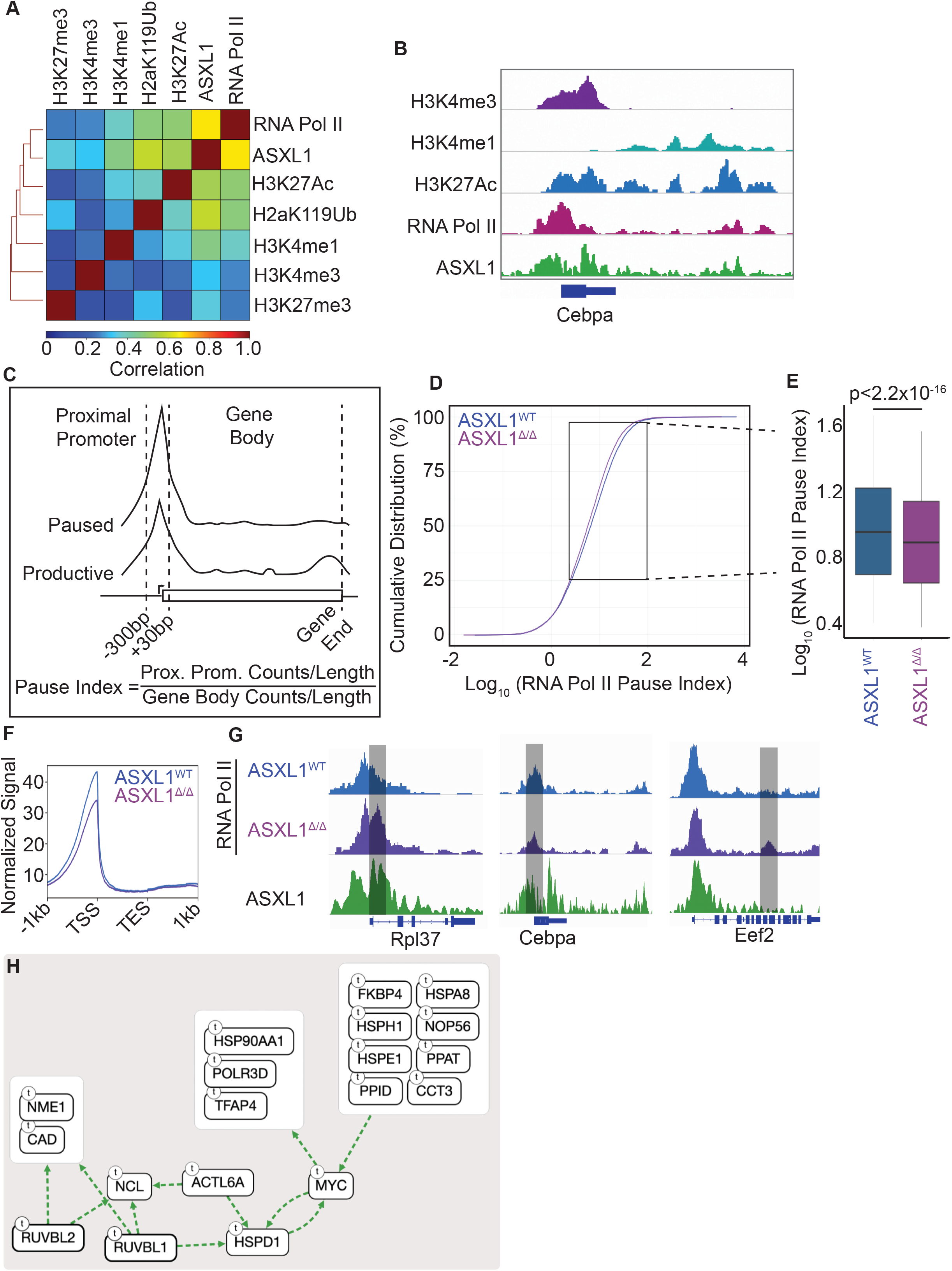
ASXL1 Deletion Leads to an Increase in RNAPII Pause-Release. **A**. Spearman correlation of histone mark or RNAPII CUT&Tag in CD133 positive granulocyte progenitors from ASXL1^WT^ mice (n=2-4/group) and published ASXL1 ChIP-seq. **B**. Representative track showing ASXL1 present across the gene body of CEBPA along with RNAPII and key covalent histone modifications. Note, H3K27me3 omitted due to a lack of signal at this actively expressed gene. **C**. Schematic depicting the calculation of RNAPII pause-index. **D**. Cumulative distribution of RNAPII pause-indices across all RNAPII bound genes from ASXL1^WT^ and ASXL1 ^Δ/Δ^ mice (n=4/group). **E**. RNAPII pause-indices within the 25-95% cumulative distribution. Statistical significance calculated by Kolmogorov-Smirnov test. **F**. Mean signal at all genes from **E. G**. Representative RNAPII tracks from genes with differential RNAPII pausing in from ASXL1^WT^ and ASXL1 ^Δ/Δ^ mice. **H**. CausalPath network analysis revealing activation of the MYC network in ASXL1-mutant AML samples from the BeatAML cohort (n=27).

### ASXL1 Mutations Lead to Disregulation of the Myc Network in Human AML

To confirm our findings in human AML samples, we examined RNA sequencing performed on 27 human ASXL1-mutant AML samples from the BeatAML cohort (Tyner et al., 2018). To examine active transcriptional networks in these samples, we performed CausalPath analysis linking genes into regulatory networks. This analysis revealed activation of a transcriptional network centered around MYC with associated activation of RNA polymerase subunits and RUVB-like family members (Figure 7H). These data confirm the activation of the MYC network in AML samples harboring ASXL1 mutations and suggests that the ASXL1-MYC-RNA Polymerase network we describe here in normal granulopoiesis may be dysregulated in human AML harboring mutant ASXL1.

## Discussion

The processes of granulopoiesis involves tightly coordinated epigenetic and transcriptional changes associated with lineage commitment and terminal differentiation (Muench et al., 2020; Olsson et al., 2016). Given that neutrophils have a very specific biological functions and peripheral blood half-life of hours to a few days, they require the expression of a relatively narrow subset of specific genes (Summers et al., 2010). Indeed, coordinated cell cycle exit and cessation of proliferation is likely necessary to prevent the malignant transformation of this high output lineage which produces an estimated one-hundred billion mature neutrophils daily (Summers et al., 2010). The proliferative balance of this lineage is disrupted in myeloid malignancy, where both over and under-proliferation are observed. In both myeloproliferative neoplasms and myelodysplastic syndromes, recurrent mutations in ASXL1 are observed, suggesting a specific connection between this gene and myeloid lineage cells (Gelsi-Boyer et al., 2012; Zhang et al., 2019). Despite extensive work on the biology of ASXL1 mutations, little is known about the role of this global regulator in normal granulopoiesis. Using single cell RNA sequencing and CUT&Tag-based low input epigenetic profiling we establish that ASXL1 plays a critical role in the regulation of active transcription, with deletion resulting in the activation of a Myc-dependent proliferative program and loss of terminal differentiation potential. These results ascribe a previously uncharacterized role to ASXL1 as a regulator of RNA polymerase activity in granulocytic lineage cells.

Numerous possible mechanisms have been proposed by which ASXL1 might mediate gene expression. Given the homeotic phenotypes associated with germline mutations, it was logically assumed that the primary role of ASXL1 in the hematopoietic system was the regulation of HOX genes (Abdel-Wahab et al., 2012; Milne et al., 1999). This was particularly compelling given the known role of HOX gene dysregulation in the pathogenesis of acute leukemia (Alharbi et al., 2013; Ayton and Cleary, 2003; Pineault et al., 2002). Indeed, in leukemia cell lines, ASXL1 deletion is associated with marked upregulation of HOXA9. However, our dataset reveals that the developmental silencing of Hox genes occurs normally in the absence of ASXL1, arguing that the disruption of terminal granulopoiesis in ASXL1^Δ/Δ^ mice occurs via a distinct mechanism. Indeed, in non-hematopoietic cell types such as bone marrow stromal cells, ASXL1 has been shown to have minimal impact on the deposition of covalent histone modifications and also similarly regulates RNA polymerase II activity (Zhang et al., 2018). Furthermore, our secondary analysis of published ChIP-seq data in HEK cells reveals a similar association of ASXL1 with regions of active transcription, arguing that this may be a broadly applicable mechanism.

The precise mechanism of ASXL1 mutations and whether they are loss or gain of function remains an area of debate, with evidence suggesting both poor expression of ASXL1 truncations as well as neomorphic interactions with BRD4 (Abdel-Wahab et al., 2012; Yang et al., 2018). Irrespective, work has largely focused on the role of ASXL1 as a regulator of covalent histone modifications. Our work suggests that the direct regulation of RNA Pol II is an important contributor to the normal function of ASXL1 in the hematopoietic system. The development of therapeutic strategies to target ASXL1 mutant clones is dependent on a comprehensive understanding of the underlying pathogenic mechanisms. Therefore, in future studies it will be critical to address the impact of mutant ASXL1 on the regulation of Myc-dependent gene expression profiles and the activity of RNA Polymerase II.

## Methods

### Mice

MX-1 Cre (003556) and ASXL1^Flox^ (025665) mice were obtained from The Jackson Laboratories. Poly I:C (Sigma) was dissolved in PBS and injected intraperitoneally at 12 mg/kg on day 1, 3 and 5 starting at 6 weeks of age. Male mice were utilized for all experiments. For in vivo GCSF treatment studies, 6-month-old ASXL1^WT^ and ASXL1 ^Δ/Δ^ mice were treated with recombinant human G-CSF (TBO-Filgrastim, Teva) at 250 μg/kg every 12 hours via intraperitoneal injection. CBCs were collected via automated blood counter (Scil Vet). All experiments were conducted in accordance with the National Institutes of Health Guide for the Care and Use of Laboratory Animals, and approved by the Institutional Animal Care and Use Committee of Oregon Health & Science University (Protocol #TR01_IP00000482).

### Flow Cytometry

The following antibodies were utilized for FACS according to the manufacturer instructions: CD150 BV421 (TC15-12F12.2, BioLegend), CD105 PE-CF594 (MJ7/18, BD), CD41a APC-Cy7 (MWReg30, Biolegend), CD16/32 PerCP e710 (93, eBioscience), Sca1 PE (D7, Biolegend), cKIT PE-Cy7 (2B8, Biolegend), Lineage APC (BD), Ly6G e450 (1A8-Ly6G, ebioscience). Stained cells were analyzed on a FACSAria III flow cytometer, LSR-Fortessa flow cytometer and Influx cell sorter (BD).

### Magnetic Cell Separation

Bone marrow cells were subjected to RBC lysis using Ammonium Chloride lysis buffer (Stem Cell Technologies), then incubated with anti-mouse direct lineage depletion cocktail (Miltenyi) according to the manufacturer’s instructions. Cells were separated using an autoMACS cell separator (Miltenyi) according to the manufacturer’s instructions. For purification of CD133 positive progenitors, bone marrow was subjected to RBC lysis as above and incubated with a biotinylated anti CD133 antibody (Clone 315-2C11, Biolegend), followed by Streptavidin MicroBeads (Miltenyi). Cells were separated using MACS LS columns (Miltenyi).

### Ex-vivo Neutrophil Generation

Lineage negative bone marrow cells were isolated as above. Cells were maintained at 3-5 X10^5^/mL throughout the culture process. Cells were initially cultured in IMDM +20 % FBS (AtlantaBiologics) in recombinant murine IL-3 and SCF (Peprotech), both at 50 ng/mL. After 3 days, recombinant murine G-CSF (Peprotech) at 50 ng/mL was added to the culture. On day 6, cells were washed 4X in PBS and resuspended IMDM +20% FBS and 50 ng/mL G-CSF. Viable cells were counted via trypan blue exclusion.

### Single Cell RNA Sequencing and CITE-seq

Lineage negative bone marrow cells (5×10^5^/mouse) were suspended in FACS staining buffer and incubated for 30 minutes with the following CITE-Seq primary antibodies and cell hash tag antibodies: CD150 (TC15-12F12.2), CD105 (MJ7/18), CD41a, CD16/32 (93), Sca1 (D7), cKIT (2B8) (BioLegend). After washing, cells were mixed in equal proportions. Mixed cell populations were loaded onto the Chromium Controller (10X Genomics) according to the manufacturer’s instructions. After the post GEM-RT cleanup, ADT and HTO additive primers were added to the RT reaction according to the manufacturer’s instructions (BioLegend). RNA and ADT/HTO libraries were separated using SPRIselect reagent and libraries were prepared separately according to the manufacturer’s instructions, Chromium Next GEM Single Cell 3ʹ v3.0 kit (Biolegend, 10X Genomics). Libraries were sequenced on a HiSeq2500 or HiSeq-X (Illumina) using 100 BP PE sequencing.

### Bulk RNA Sequencing

Total RNA was extracted from approximately 5×10^5^ CD133 purified progenitors using an RNeasy micro kit (Qiagen) according to the manufacturer’s instructions. Libraries were prepared using a TruSeq library prep kit (Illumina) according to the manufacturer’s instructions. Libraries were sequenced on a HiSeq2500 (Illumina) using SE 100BP sequencing.

### Protein-A Tn5 Transposase Preparation

The pA-Tn5 plasmid was a generous gift from Steven Henikoff (Addgene Plasmid #124601). The plasmid was maintained in T7 Express lysY/Ig Competent *E.coli* cells (NEB). To prepare the transposase, a 3 mL LB starter culture with 100 μg/mL carbenicillin (Sigma) was prepared from a single fresh colony and grown for 4 hours at 37° C. This culture was then used to inoculate a 400 mL LB culture with 100 μg/mL carbenicillin (Sigma) and grown until an OD_600_ of 0.5 to 0.6 was reached. The culture was then transferred to 4° C for 1 hour, after which IPTG was added to a final concentration of 0.25mM. The culture was then shaken for 14 hours at 18°C. The culture was then spun down at 10,000 RPM for 30 minutes at 4°C and the pellet was frozen in a dry ice/ethanol bath and stored at −80°C. The pellet was then resuspended in HEGX Buffer (20 mM HEPES-KOH, pH 7.2, 0.8M NaCl, 1 mM EDTA, 10% glycerol, 0.2% Triton X-100 and 1 Roche Complete Protease Inhibitor Tablet, EDTA free in 50 mL buffer). After resuspension, the pellet was sonicated using a Digital Sonifier (Branson) equipped with a 2 mm tip for 10 cycles of 45 seconds with a 50% duty cycle. Lysate was kept on ice for the duration of the sonication. After sonication, the lysate was centrifuged at 10,000 RPM for 30 minutes at 4° C, and the supernatant was loaded onto two disposable chromatography columns (BioRad) loaded with 2.5 mL chitin slurry resin (New England Biolabs). After loading, columns were rotated at 4°C overnight. Columns were then drained and washed 2X with 20 mL of HEGX buffer. To elute the protein, 6 mL HEGX buffer supplemented with 100 mM DTT was added to each column, and the columns were rotated at 4°C for 48 hours. The columns were then drained and the eluate was dialyzed against 1 L of 2X Tn5 dialysis buffer (100 mM HEPES-H pH 7.2, 0.2 M NaCl, 0.2 mM EDTA, 2 mM DTT, 0.2% Triton X-100, 20% glycerol) for 2 hours and then again overnight using Slide-A-Lyzer Dialysis Cassettes (Thermo). Eluate was then concentrated to 300 uL using a Centrifugal Filter Unit with 30 kDa size cutoff (Sigma) by centrifugation at 4000 RPM at 4°C. The concentrated protein was diluted to 50% glycerol, quantified by comparison to BSA standards using an SDS Page Gel (BioRad).

### CUT&Tag

ME-A and ME-B adaptors were annealed separately with the ME-R adaptor and then mixed (Kaya-Okur et al., 2019). The mixed adaptors (16 μL of 100 μM stock) were then added to 100 uL of ~5.5 μM pA-Tn5 transposase. The mixture was rotated at room temperature for 1 hour then stored at −20°C prior to use. Sorted or purified cells (1×10^4^ to 1×10^5^) were washed in CUT&Tag wash buffer 2X (20 mM HEPES pH 7.5, 150 mM NaCl, 0.5 mM Spermidine, 1× Protease inhibitor cocktail). Concanavlin A magnetic coated beads (Bangs Laboratories) were activated by washing in binding buffer (20 mM HEPES, pH 7.5, 10 mM KCL, 1 mM CaCl_2_, 1 mM MnCl_2_) 2X. To bind the cells to the beads, 10 uL of beads were added to each sample and rotated end over end for 7 minutes. Using a magnetic stand, supernatant was removed and primary antibody was added to each sample at a 1:50 dilution in antibody buffer (20 mM HEPES pH 7.5, 150mM NaCl, 0.5 mM Spermidine, 1× Protease inhibitor cocktail, 0.05% digitonin, 2 mM EDTA, 0.1% BSA). The following primary antibodies were utilized: H3K27Ac (ab4729, Abcam), H3K4me1 (#5326, Cell Signaling Technologies), H3K4me3 (#9751, Cell Signaling Technologies), H3K27me3 (#9733, Cell Signaling Technologies), H2aK119Ub (#8240, Cell Signaling Technologies). Cells were incubated on a nutator at 4°C overnight, then antibody was removed and a guinea-pig anti rabbit secondary antibody was added at 1:100 (Antibodies Online). Samples were incubated on a nutator at room temperature for 1 hour. Secondary antibody was removed and the samples were washed twice in digitonin-wash buffer (20 mM HEPES pH 7.5, 150 mM NaCl, 0.5 mM Spermidine, 1× Protease inhibitor cocktail, 0.05% digitonin). pA-Tn5 transposase was diluted 1:250 in digitonin-300 buffer (20 mM HEPES pH 7.5, 300 mM NaCl, 0.5 mM Spermidine, 1× Protease inhibitor cocktail, 0.01% digitonin) and samples were incubated on the nutator for 45 minutes at room temperature. Samples were then washed twice with digitonin-300 buffer and then resuspended in tagmentation buffer (digitonin 300 buffer supplemented with 1 mM MgCl_2_) and incubated at 37°C for 1 hour. DNA was then extracted by phenol:chloroform extraction. Samples were then amplified by PCR using custom nextera primers at 400 nM and NEBNext HiFi 2x PCR Master Mix (New England Biolabs) (Buenrostro et al., 2017). PCR conditions were as follows: 72°C for 5 minutes; 98°C for 30 seconds; 14 cycles of 98°C for 10 sec, 63°C for 10 sec; and 72°C for 1 minute. Libraries were purified with AMPure Beads (Beckman) and sequenced on a NextSeq 550 sequencer (Illumina) using 37 PE sequencing.

### Data Analysis

#### Single Cell RNA Sequencing Analysis

Sequencing output from 10X mRNA, ADT and HTO sequencing libraries were aligned to the murine genome (mm10) using CellRanger. Filtered feature matrices were analyzed using Seurat (Butler et al., 2018). Cells expressing greater than 20% mitochondrial RNA were excluded as non-viable. Cells were assigned to individual mice via labeling with a specific HTO barcode. Cells labeling with two distinct HTO barcodes were excluded as doublets. Pseudo-flow cytometry plots were generated using Log normalized CITE-seq antibody read counts and gates were drawn according to published descriptions (Oguro et al., 2013; Pronk et al., 2007). Data integration revealed 25 transcriptional clusters. Cluster identify was established via correlation with published murine RNA seq from sorted cell populations (Choi et al., 2019) coupled with manual curation. Differential expression by cluster was performed by Wilcox-based analysis using Seurat and GO analysis performed using TopGO from Bioconductor. Lineage trajectories were identified using Monocle3 (Qiu et al., 2017a, 2017b; Trapnell et al., 2014). CytoTRACE analysis was performed as previously reported using expression matrices and the CytoTRACE R package (Gulati et al., 2020)

#### Causal Path analysis

Causal path statistically evaluates and mechanistically grounds relationships observed among transcriptomic, proteomic and phosphoproteomic measurements to yield inferences of activity flow across data points. This method is explained in more detail in a preprint under the name CausalPath (Babur et al., 2018). CausalPath is a hypothesis generation tool that automates the literature search of a biologist. This method aids researchers in understanding experimental observations using known mechanisms with a focus on post-translational modifications. Experimental data reveal protein features that change in coordination, and CausalPath automates the search for causal explanations in the literature. The identified relations are not certainty, but rather hypotheses that can guide discovery. CausalPath generates causal explanations for the observed correlations in the dataset using literature-curated pathway data. It relies on a library of graphical patterns that represent a variety of possible relationships occurring between pairs of proteins. CausalPath pares down the changes observed in the (phospho)proteomic or transcriptomic data to include only those which are both strongly correlated and causally explainable by this library. These explainable relationships are then reduced to an intuitive, simplified network which can be graphically represented via the ChiBE desktop visualization suite (Babur et al., 2018)

#### Bulk RNA Sequencing Analysis

Raw reads were trimmed with Trimmomatic (Bolger et al., 2014) and aligned with STAR (Dobin et al., 2013). Differential expression analysis was performed using DESeq2 (Love et al., 2014). Raw p values were adjusted for multiple comparisons using the Benjamini-Hochberg method. GO analysis was performed using Enrichr (Kuleshov et al., 2016).

#### CUT&Tag Analysis

CUT&Tag libraries were aligned to the mouse genome (mm10) using Bowtie2 (Langmead and Salzberg, 2012) and the following options --local --very-sensitive-local --no-unal --nomixed --no-discordant -- phred33 -I 10 -X 700. BAM files were downsampled to the lowest common denominator of read depth across conditions using Samtools (Li et al., 2009). To avoid issues with false positives inherent to CUT&Tag data, peaks were called on a merged BAM file containing equal representation from all conditions using MACS2 (Zhang et al., 2008). A merged IgG negative control BAM was used as the control. The following q-value cutoffs were used for each specific histone mark to ensure capture of high confidence peaks only: H3K4me1 q= 0.00001; H3K4me3 q= 0.0001 for peaks, q= 1×10^−26^ to exclude low abundance peaks at enhancers; H3K27Ac q=0.001, H3K27me3 q=0.001, H2aK119Ub q= 0.000001, Rbp1 q=0.01. Differential peaks were identified utilizing the Bioconductor package Diffbind using the default parameters (Ross-Innes et al., 2012). Heatmaps were produced using the ComplexHeatmap package from Bioconductor (Gu et al., 2016). Peak annotation and GO analysis were performed using ChIPseeker (Yu et al., 2015). To identify RNAPolII bound genes, all mm10 genes were downloaded from the UCSC table browser and intersected with Rbp1 peaks using Bedtools (Quinlan and Hall, 2010). Counts tables for all analyses were generated using featureCounts from the subread package (Liao et al., 2013). Global signal correlation and heatmaps were generated using DeepTools (Ramírez et al., 2016). RNAPolII pause indices were calculating by generating counts tables for each gene in a window from −30 to +100 around each TSS and then from the remainder of the gene bodies. Counts were divided by the region length and the pause index was calculated by dividing the TSS values by those for the corresponding gene body. Published ChIP-seq data was downloaded from Cistrome DB with the following accession numbers: ASXL1 76682, H3K4me1 88060, H3K4me3 76226, H3K27Ac 66607, Rbp1 91539 (Fong et al., 2017; Hauri et al., 2016; Morgan et al., 2017; Savitsky et al., 2016).

## Acknowledgements

We would like to thank Dr. Marilynn Chow-Castro for her thoughtful discussions; the OHSU Massively Parallel Sequencing Shared Resource for scRNA-seq library prep using their 10X Genomics Chromium Controller and performing short read sequencing assays. This project was supported by funding from the Cancer Early Detection and Research center (CEDAR) at Oregon Health and Science University’s Knight Cancer Institute.

---

**Supplementary Figure 1.**
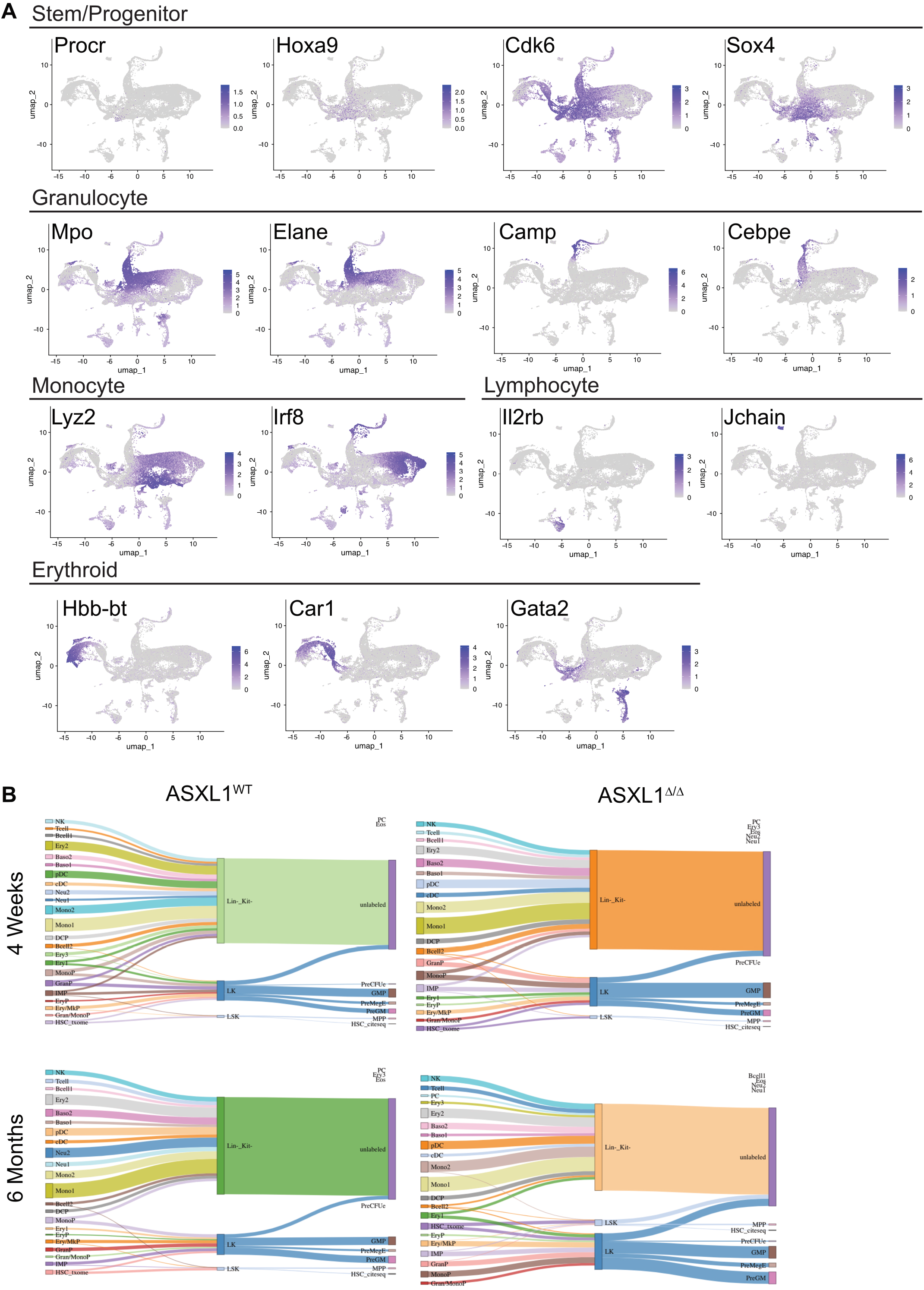
Combined Transcriptional and Proteomic Signatures Define Bone Marrow Lineages. **A**. Feature Plots of key lineage defining marker genes. **B**. Sankey plots showing the relationship between transcriptional and CITE-seq defined clusters.

**Supplementary Figure 2.**
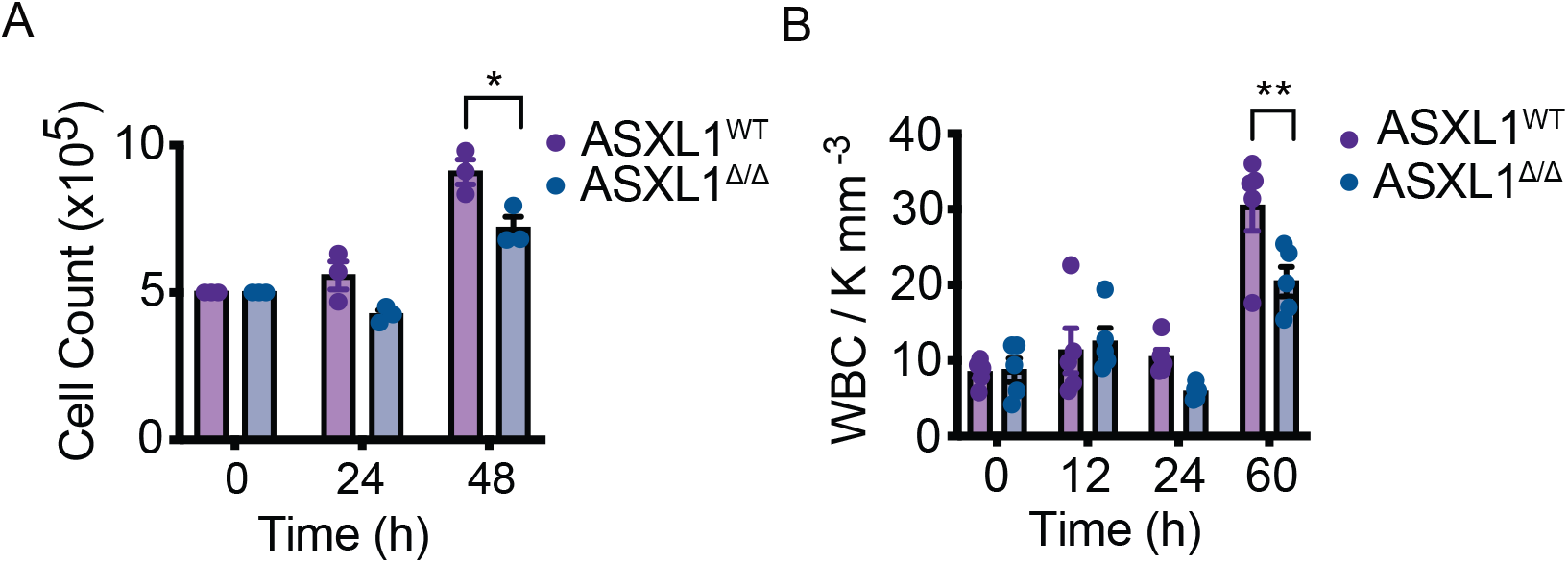
ASXL1 Deletion Leads to Impaired Neutrophil Production in vitro and in vivo. **A**. Lineage negative cells were isolated from the bone marrow of ASXL1^WT^ and ASXL1 ^Δ/Δ^ mice (n=3/group) and grown for 3 days in IL-3+SCF, then 3 days in IL-3+SCF+G-CSF. At this time cell number was normalized between mice and cells were cultured for 2 days in G-CSF only. Full neutrophilic differentiation was confirmed after 48 hours and was equivalent between genotypes. **B**. ASXL1^WT^ and ASXL1 ^Δ/Δ^ mice (n=5/group) were treated with G-CSF 250 μg/mL) every 12 hours. WBC counts were collected at the indicated time points. In all cases, data is represented as the mean +/− SEM. * = p<0.05, ** = p<0.01 by two-way ANOVA with Sidak’s post-test.

**Supplementary Figure 3.**
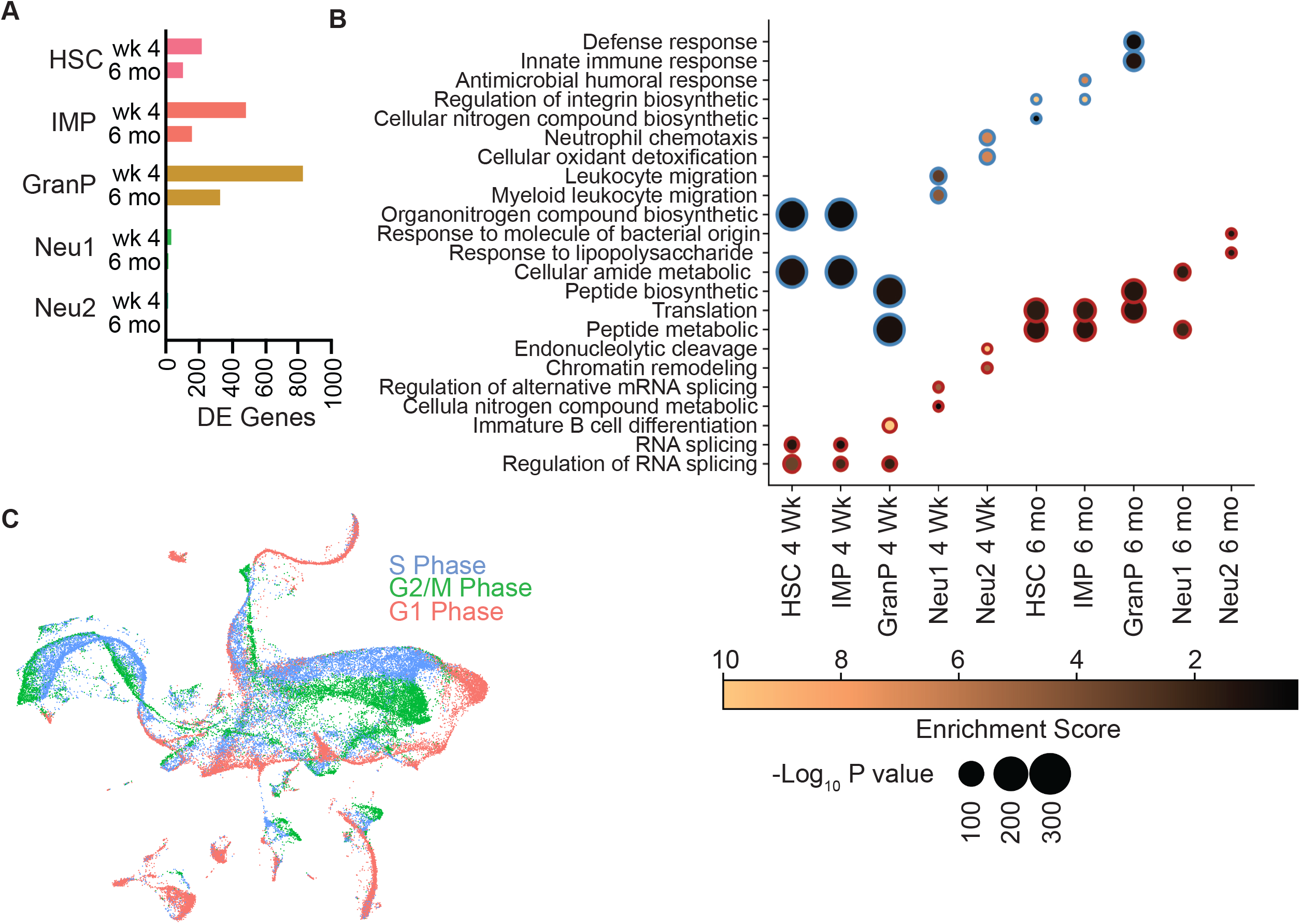
ASXL1 Deletion Produces Maximal Transcriptional Dysregulation in Granulocyte Progenitors. **A**. Number of differentially expressed genes in transcriptionally defined clusters comprising the granulocyte lineage in ASXL1^WT^ and ASXL1 ^Δ/Δ^ mice from Figure 3. **B**. Gene ontology analysis performed on differentially expressed genes from **A** and Figure 3. **C**. UMAP projection demonstrating that exit from the granulocyte progenitor is associated with cell cycle arrest.

**Supplementary Figure 4.**
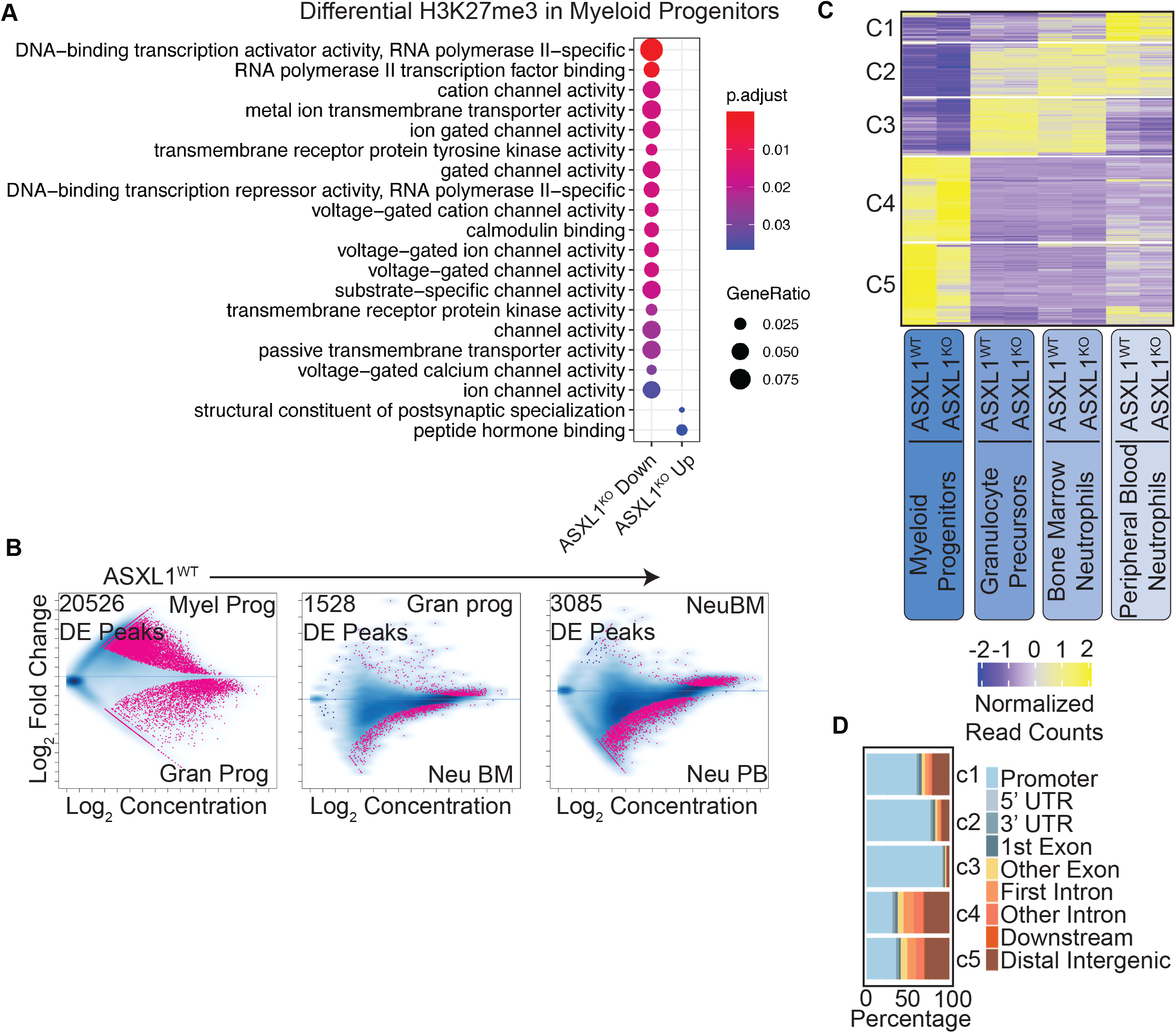
ASXL1 Deletion Alters H3K27me3 Deposition in Bipotent Myeloid but not Unipotent Granulocytic Progenitors. **A**. GO analysis on regions of differential H3K27me3 enrichment in myeloid progenitors. **B**. Differential enrichment analysis of H3K27me3 in ASXL1^WT^ mice between myeloid progenitors, granulocyte progenitors, bone marrow neutrophils and peripheral blood neutrophils. **C**. Row normalized heatmap of H3K27me3 counts in a differential regions catalog assembled from the comparisons in **B** and in Figure 5C. Counts are row normalized to highlight genotype and cells state differences rather than changes in absolute mark abundance. Clustering of rows performed by K-means. **D**. Genome wide localization of clusters from **C**.

**Supplementary Figure 5.**
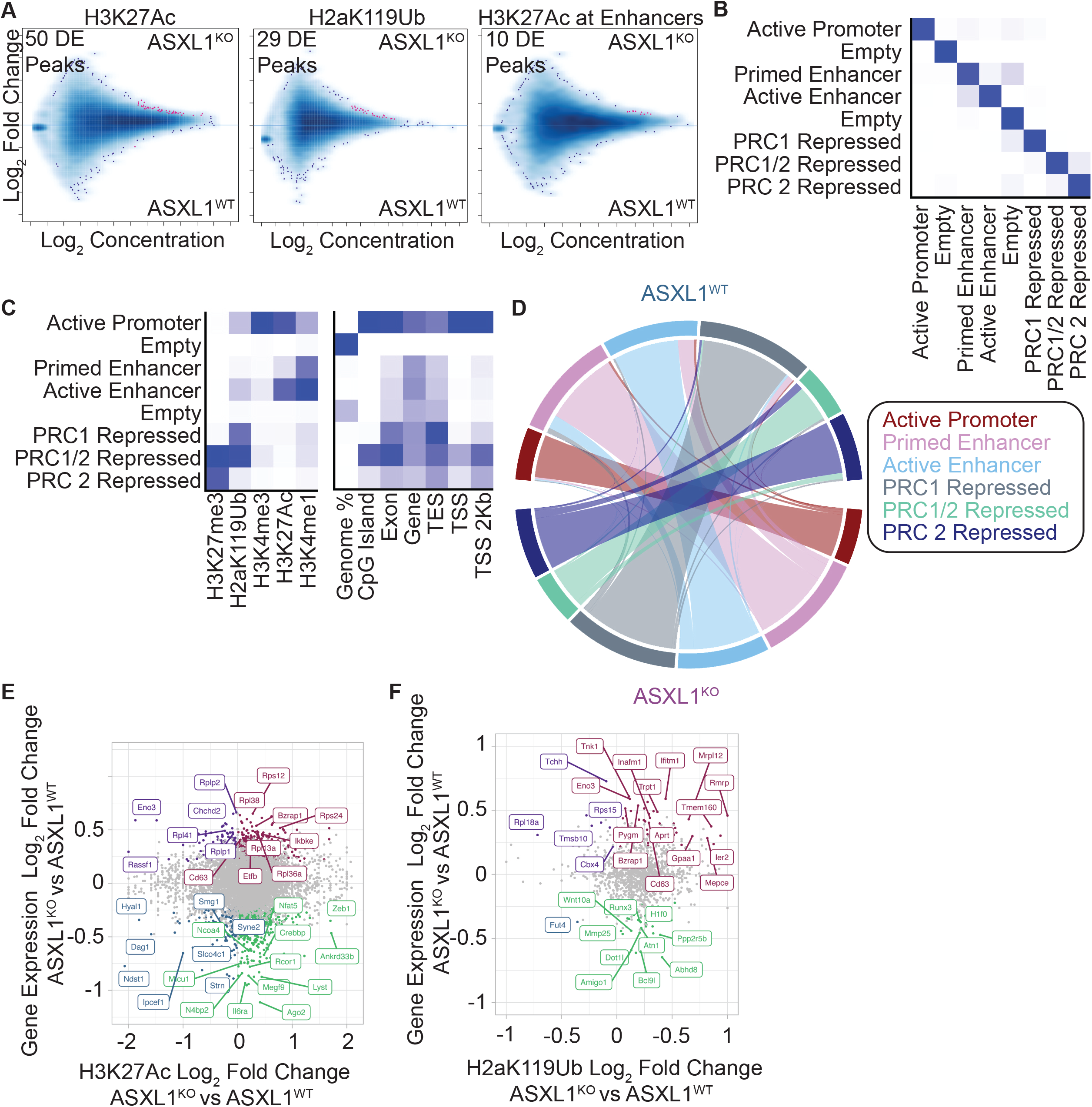
Chromatin Changes in Response to ASXL1 Deletion in Granulocytic Progenitors. **A**. Differential enrichment analysis on CUT&Tag for indicated histone marks from Figure 6E, F. **B**. Transition matrix from chromatin segmentation analysis using CUT&Tag for the indicated histone marks from Figure 6 E, F using ChromHMM. **C**. Emission matrix demonstrating co-localization of covalent histone modifications and their genome wide localization. **D**. Chord diagram showing chromatin state changes that occur in granulocyte progenitors upon deletion of ASXL1. **E**. Correlation between H3K27Ac fold change and gene expression changes observed in nearest gene by bulk RNA seq from Figure 6C. **F**. Correlation between H2aK119Ub fold change and gene expression changes observed in nearest gene by bulk RNA seq from Figure 6C.

**Supplementary Figure 6.**
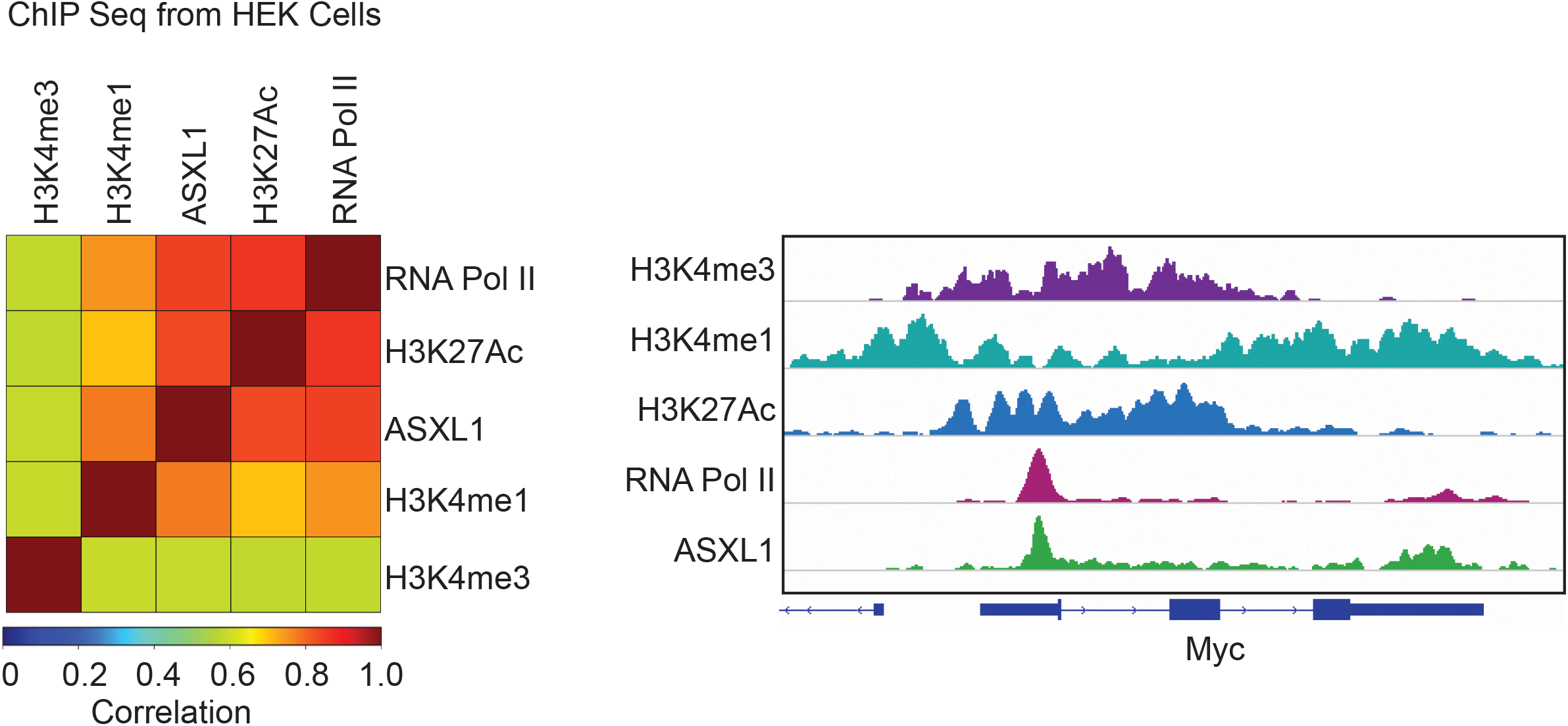
Co-Localization of ASXL1 with RNAPII in HEK Cells. Pearson correlation of histone mark or RNAPII CUT&Tag in HEK293 from published data. Representative track showing ASXL1 present across the gene body of MYC along with RNAPII and key covalent histone modifications. Note, H3K27me3 omitted due to a lack of signal at this actively expressed gene.

